# AraENCODE: a comprehensive epigenomic database of *Arabidopsis Thaliana*

**DOI:** 10.1101/2023.06.10.544382

**Authors:** Zhenji Wang, Minghao Liu, Fuming Lai, Qiangqiang Fu, Liang Xie, Yaping Fang, Qiangwei Zhou, Guoliang Li

**Author notes:** Correspondence: Guoliang Li. These authors contributed equally to this article (Zhenji Wang, Minghao Liu).

## Abstract

Arabidopsis (*Arabidopsis thaliana*) is a vital model organism in plant biology and genetics. In the last two decades, researchers have made significant progresses in characterizing the chromatin conformation and epigenomic information within the Arabidopsis genome. This information includes but is not limited to the higher structure of chromosomes, histone modification, DNA methylation, and chromatin accessibility. The results of these studies have provided an additional layer of information that complements the DNA sequence data. However, utilizing such knowledge poses a challenge for certain groups that lack bioinformatics analysts or adequate computing resources. A user-friendly and reproducible platform for accessing this information is urgently needed. In this study, we have developed a comprehensive epigenomic database for Arabidopsis (AraENCODE http://glab.hzau.edu.cn/AraENCODE), which comprises a total of 4,511 data libraries, including published chromatin conformation capture datasets (Hi-C, HiChIP), epigenomic datasets (ChIP-Seq, ATAC-Seq, FAIRE-Seq, MNase-Seq, DNase-Seq, BS-seq), and transcriptome data (RNA-Seq, miRNA-Seq). Furthermore, we have incorporated various existing resources, such as single nucleotide polymorphisms (SNPs), cis-regulatory modules, and multi-omics associations. We aim to provide a novel platform for investigating the regulation of epigenetic and chromatin interactions in Arabidopsis in relation to biological processes.

## Main text

Arabidopsis (*Arabidopsis thaliana*) is an important model organism in plant biology and genetics. The genome of Arabidopsis ecotype Columbia-0 (Col-0) has been sequenced and completely annotated (Cheng et al., 2017), which facilitates the genomics research of plants. Over the past two decades, advances in the gene regulation study have elucidated a spectrum of epigenetic molecular phenomena, including DNA methylation, histone modification, chromatin accessibility, and chromatin interaction, which collectively form an additional layer of information based on DNA sequence (Law and Jacobsen, 2010). The epigenome landscape has been characterized in Arabidopsis as more high-throughput analyses were developed (Zhao et al., 2022). Without a doubt, in-depth studies of gene expression regulation heavily rely on such epigenomic information. However, the utilization of this knowledge poses a challenge for certain groups lacking bioinformatics analysts or adequate computing resources.

Here, we developed a comprehensive epigenomic database for Arabidopsis (AraENCODE, http://glab.hzau.edu.cn/AraENCODE/), which comprises a total of 4,511 sample accessions from Sequence Read Archive (SRA), Gene Expression Omnibus (GEO), Genome Sequence Archive (GSA) and other open-access databases, including datasets with histone modification, chromatin accessibility, DNA methylation, transcriptome, and chromatin interactions from different tissues in wild type or mutants (Figure 1A). The resource and distribution of the data sets are displayed in detail on the “Data Statistics” module of the website (http://glab.hzau.edu.cn/AraENCODE/pages/datasets.html) (Supplemental Figure 1). We downloaded raw data from various libraries encompassing ChIP-seq, ATAC-seq, DNase-seq, MNase-seq, FAIRE-Seq, Hi-C, HiChIP, BS-seq, RNA-seq, and ncRNA-seq (Supplemental Table 1), and subsequently reprocessed them using a standardized pipeline tailored to each data type (Figure 1B). Quality control metrics for various types of data have been provided to identify the data quality in the “Quality Control” module. For the sake of facilitating comparisons between various datasets, all datasets were aligned to the TAIR10 genome and visualized using the WashU epigenome browser (Li et al., 2022), which can help the users compare different tracks flexibly. Such a database provides a comprehensive view of the relationships between gene expression and the epigenetic landscape across various tissues and genotypes in Arabidopsis. Furthermore, it has the potential to facilitate our understanding of the epigenetic mechanism in higher plants.

**Figure 1.**
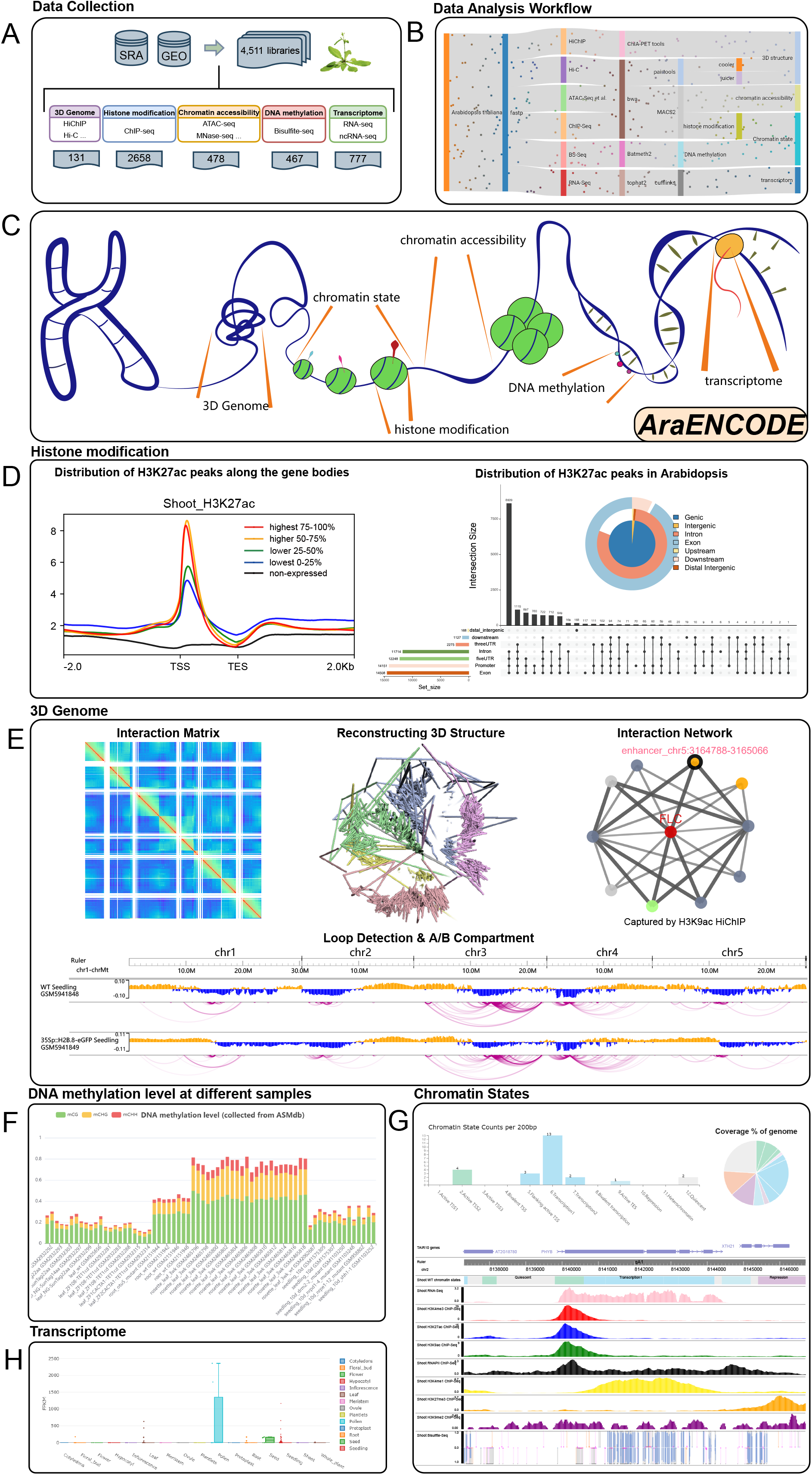
Architecture and screenshots of the AraENCODE database. (A) Data collection of AraENCODE. (B) A standardized pipeline for different data types (Code accessibility: https://github.com/versarchey/AraENCODE-pipeline). (C) AraENCODE website (http://glab.hzau.edu.cn/AraENCODE). (D) The “Histone Modification” module shows modification peaks in different samples and their distribution over the whole genome. Here is an example of H3K27ac in shoot tissue. (E) The “3D Genome” module shows chromatin interactions in different samples. Interactions matrixes are visualized by HiGlass browser and are used to reconstruct 3D structures, detect loops and make compartment analysis. Interaction network constructed by loops shows the gene-gene or gene-enhancer interactions. This is an example of network involved gene AT5G10140 (also known as FLC or RSB6). (F) DNA methylation level at different samples. (G) Example of chromatin states search results (around gene AT2G18790, also known as PHYB). (H) Example of gene expression level among different tissues (pollen-specific expression gene AT5G45880, also known as LAT52).

According to various functions, AraENCODE is divided into the following parts: “Quick Search”, “WashU Browser”, “Histone Modification”, “3D Genome”, “Open Chromatin”, “Chromatin State”, “DNA Methylation”, “Transcriptome”, “WT/Mutant”, “Datasets”, “Download”, “Analysis Pipeline” and “Tutorial”. In AraENCODE (Figure 1C), users can retrieve epigenomic information by querying a gene ID or a genome region. On the home page, users could query a gene ID and fetch all levels of epigenome information related to this gene simultaneously (Supplemental Figure 2A). The quick search results are intended to provide researchers with a concise overview of the epigenomics information pertaining to the gene of interest. The quick search page contains a WashU browser window that includes a few tracks: the 7 types of histone modifications and DNA methylation of the Arabidopsis genome as well as chromatin states and SNP information (Supplemental Figure 2B); Specific protein-mediated chromatin loops which link regulatory elements (e.g., enhancers) or single nucleotide polymorphisms (SNPs) physically close to their target genes, and can be characterized using 3C-based methods (e.g., Hi-C and HiChIP), histone modification, and chromatin accessibility information (Supplemental Figure 2C); Profiles of cross-tissue differential expression and differential methylation levels (Supplemental Figure 2D, E). For more detailed information, users have the option to navigate to the corresponding page for in-depth exploration.

Histones are often covalently modified to affect various chromatin-dependent processes, including gene transcription. Within the histone modification page, users can search for genes or chromosome regions and retrieve information regarding the various histone modification and their distribution over the whole genome in different samples (Figure 1D). AraENCODE also provides information on chromatin accessibility, which is important for establishing and maintaining cell identity and can be characterized by ATAC-Seq, FAIRE-Seq, DNase-Seq and MNase-Seq. This information can be viewed in a table or visualized through the WashU epigenome browser. As a validation, we examined the abundance of histone modification on well-documented genes in our database, such as flowering time regulation gene AT5G10140 (also known as FLC or RSB6). Previous studies have indicated a correlation between the overexpression of COLDAIR (lncRNA) in mutant 35S-COLDAIR and the down-regulation of the repressive histone mark H3K27me3, as well as the up-regulation of the active histone mark H3K4me3 at FLC gene, which upregulates the FLC expression and ultimately impacts the flowering process (Liu et al., 2020). The abundance of these histone modifications and gene expressions in our database is highly consistent with previous reports (Supplemental Figure 3).

DNA methylation (5-methylcytosine), a stable epigenetic mark, underpins the landscape of histone modifications in Arabidopsis (Zhao et al., 2022). Three sequence contexts of DNA methylation (CG, CHG, and CHH) are catalyzed by different methyltransferases and interdependently occur in different contexts (Law and Jacobsen, 2010). On the “DNA Methylation” page, users can quickly browse the methylation levels across tissues (Figure 1F). The gene search module enables users to retrieve DNA methylation levels of three sequence contexts (CG, CHG, and CHH) in the gene body or promotor regions (Supplemental Figure 4A-D), and browse it at a single-base resolution across all samples (Supplemental Figure 4E) (Zhou et al., 2022). As an example, we investigated the expression levels and methylation levels of two genes AT3G50770 (CML4) and AT5G43260 in three mutants, including met1, ddcc (drm1 drm2 cmt2 cmt3), and mddcc (met1 drm1 drm2 cmt2 cmt3), with different degrees of demethylation, as well as in the wild type (Col-0). We found that the expression of AT3G50770 is up-regulated in the mutants (Supplemental Figure 5A), especially in the mddcc mutant whose DNA methylation in all contexts is eliminated. Meanwhile, the expression of AT5G43260 does not change significantly (Supplemental Figure 5B). Such results are consistent with previous reports (Naydenov et al., 2015).

Chromatin states, which reflect genome activity and transcriptional regulation in eukaryotes (Roudier et al., 2011), are mainly determined by histone modification and DNA methylation. To train the model and achieve the precise classification of chromatin states (Zhao et al., 2022), we utilized ChromHMM along with six types of modifications, RNAPII occupancy, and methylation across diverse varieties. The Arabidopsis genome can be segmented into 12 chromatin states with distinct percentages of genome coverage. On the chromatin states page, users could select different varieties and search a gene or a region to obtain the chromatin state information at a resolution of 200 base pairs (bp) (Figure 1H).

Previous studies have shown that gene expression can be regulated by the three-dimensional structure of chromatin (Zhang et al., 2019). In AraENCODE, we collected the Hi-C, Capture Hi-C and HiChIP datasets to build the “3D Genome” to shows chromatin interactions in different samples. Interactions matrixes from Hi-C and Capture Hi-C are visualized using the HiGlass browser, enabling the examination of global variations among different samples. Hi-C and Capture Hi-C are also used to reconstruct 3D structures, detect chromatin loops and make compartment analysis. These results can be checked in the WashU browser.

HiChIP has been developed to detect and quantify chromatin contacts anchored at genomic regions associated with specific DNA binding proteins or histone modifications, similar to ChIA-PET (Fullwood et al., 2009; Mumbach et al., 2016). The HiChIP workflow includes the following steps: cell lysis and permeabilization, in-situ restriction enzyme digestion, biotin labeling and in-situ proximity ligation, chromatin immunoprecipitation (chromatin shearing, IP and wash) and library preparation. By integrating in-situ Hi-C and chromatin immunoprecipitation (ChIP), HiChIP is able to detect long-range chromatin contacts mediated or associated with specific protein factors at kilobase-scale resolution with significantly reduced sequencing cost compared to in-situ Hi-C (Rao et al., 2014). With HiChIP in Arabidopsis, Huang and colleagues show that H3K27me3 is a key regulator of global and local facultative heterochromatin topology, and is tightly linked to 3D organization (Huang et al., 2021).

To detect significant chromatin interactions, HiChIP datasets in AraENCODE were reprocessed using ChIA-PET Tool (V3), a computational package designed for processing sequence data derived from ChIA-PET or HiChIP experiments (Li et al., 2019). Interaction networks, constructed through chromatin loops, reveal gene-gene or gene-enhancer interactions (an example that involved gene AT5G10140, also known as FLC or RSB6, are shown). In “3D Genome” module, users can query a gene or region, especially phenotype-associated GWAS SNPs, to find its target loci and visualize these interactions using the built-in browser (Figure 1E). And an interaction network is provided for users to examine the loops connecting the genes and regulatory elements (Figure 1E). We investigated the SNPs associated with AT5G10140 (FLC) expression in AraGWAS (1,241 SNPs in total, 623 within the FLC gene body and 618 outside of FLC gene body) (Supplemental Table 2). 91% (563/618) of the SNPs outside of FLC gene body are located within the anchors (PET count >= 10) interacting with FLC (Togninalli et al., 2020), which includes a lot of SNPs with high scores in AraGWAS (e.g., chr5:3181599 G>C and chr5:3181811 C>A). These SNPs that overlapped with the interaction regions were concentrated in two regions (chr5:3168751-3174117 and chr5:3180175-3189848), involving the genes AT5G10120, AT5G10130 and AT5G10150, and the detailed interaction network are also provided for a more comprehensive view in Supplemental Figure 6. These findings suggest a potential mechanism that these extragenic SNPs contribute to the regulation of FLC expression. To our knowledge, this is the most comprehensive resource for Arabidopsis 3D genome data to date.

AraENCODE also collected high-throughput transcriptomic data from various tissues. AraENCODE includes expression levels of specific genes as well as miRNA (Figure 1H). Although there are existing RNA-seq databases dedicated to Arabidopsis, such as ARS (Zhang et al., 2020), we firmly believe that incorporating RNA-seq data into our database will provide users with a more convenient means of integrating transcriptomic and epigenomic information for further analysis.

As Arabidopsis high-throughput data accumulate, many excellent web-based resources have been developed. For example, the AraGWAS Catalog (Togninalli et al., 2020) focuses on genotype-phenotype associations, ARS (Zhang et al., 2020) collects high-throughput transcriptome data, and the ChIP-Hub focuses on regulome data of plants (Fu et al., 2022). However, to our knowledge, AraENCODE is the first comprehensive, user-friendly, and sustainably maintained database for epigenome and 3D genome in Arabidopsis. AraENCODE provides a comprehensive map of the Arabidopsis epigenetic landscape, facilitating rapid access to the epigenomic information, chromatin states, and 3D interactions associated with GWAS SNPs and their target genes, which could further improve our knowledge of gene regulation mechanisms. In future, with the rapid generation of high-throughput data, we will regularly integrate additional resources and valuable analytical tools for Arabidopsis epigenomics and 3D genomics.

## Acknowledgments

This work was supported by the Fundamental Research Funds for the Central Universities (2662021PY005 to G.L.).

## Code availability

The source code of the data analysis pipeline is available from the GitHub repository (https://github.com/versarchey/AraENCODE-pipeline).

## Author contributions

G.L. conceived and supervised the project. Z.W. and M.L. designed the database structure; M.L. collected datasets and analyzed the data; F.L. and Q.F. collected Hi-C datasets and analyzed the data. L.X. collected some datasets and helped in data analysis; Q.Z. provided some of the analysis results; Z.W. collated analysis results, built the database, and drafted the initial manuscript. All authors read, revised, and approved the final manuscript.

