## Supplemental Figure for "AraENCODE: a comprehensive epigenomic database of *Arabidopsis Thaliana*"

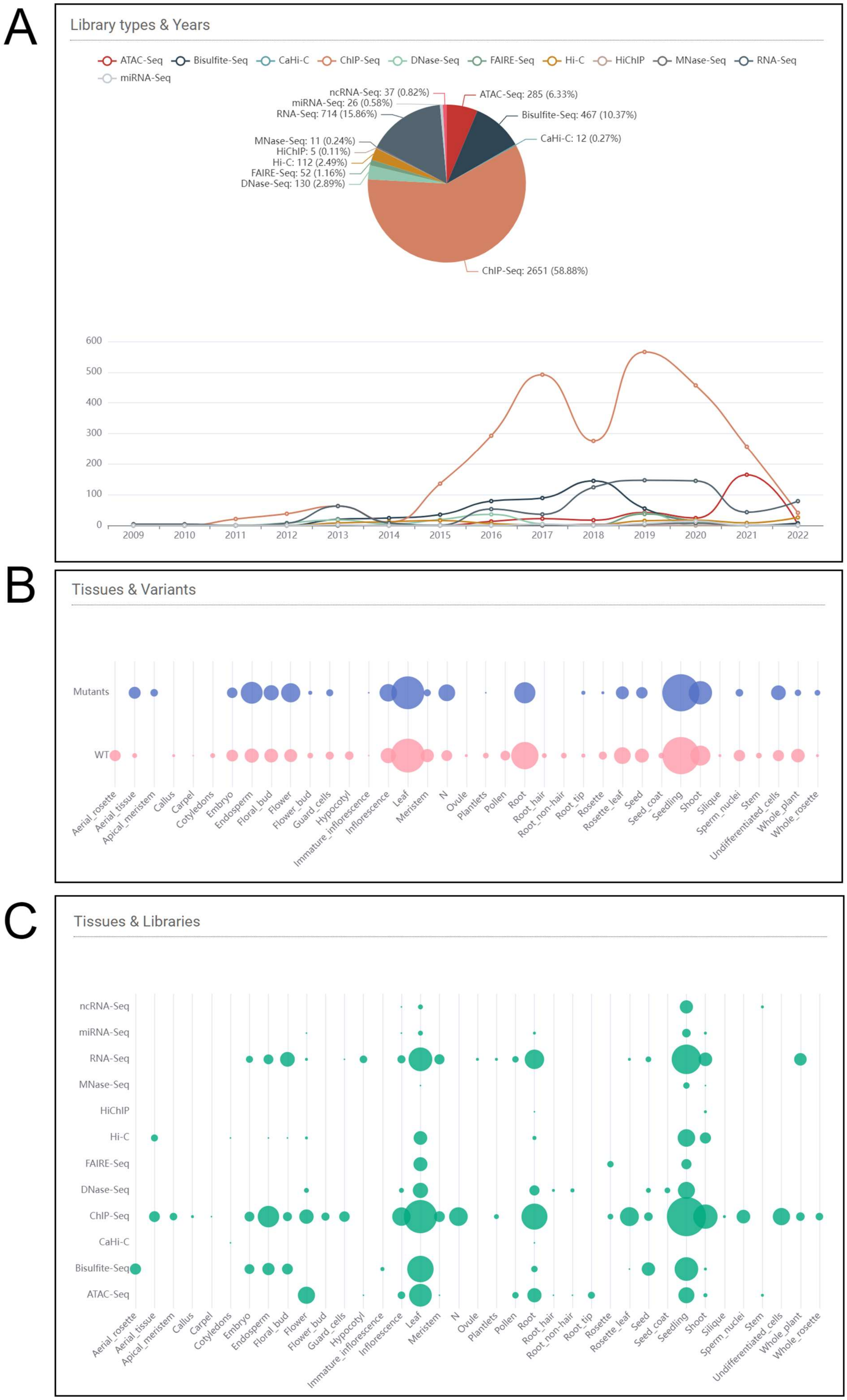
A
Library types & Years
ATAC-Seq
Bisulfite-Seq
CaHi-C
ChIP-Seq
DNase-Seq
FAIRE-Seq
Hi-C
HiChIP
MNase-Seq
RNA-Seq
miRNA-Seq
ncRNA-Seq: 37 (0.82%)
miRNA-Seq: 26 (0.58%)
RNA-Seq: 714 (15.86%)
ATAC-Seq: 285 (6.33%)
Bisulfite-Seq: 467 (10.37%)
CaHi-C: 12 (0.27%)
MNase-Seq: 11 (0.24%)
HiChIP: 5 (0.11%)
Hi-C: 112 (2.49%)
FAIRE-Seq: 52 (1.16%)
DNase-Seq: 130 (2.89%)
ChIP-Seq: 2651 (58.88%)
600
500
400
300
200
100
0
2009
2010
2011
2012
2013
2014
2015
2016
2017
2018
2019
2020
2021
2022
B
Tissues & Variants
Mutants
WT
Aerial_rosette
Aerial_tissue
Apical_meristem
Callus
Carpel
Cotyledons
Embryo
Endosperm
Floral_bud
Flower
Flower_bud
Guard_cells
Hypocotyl
Immature_inflorescence
Inflorescence
Leaf
Meristem
N
Ovule
Plantlets
Pollen
Root
Root_hair
Root_non-hair
Root_tip
Rosette
Rosette_leaf
Seed
Seed_coat
Seedling
Shoot
Silique
Sperm_nuclei
Stem
Undifferentiated_cells
Whole_plant
Whole_rosette
C
Tissues & Libraries
ncRNA-Seq
miRNA-Seq
RNA-Seq
MNase-Seq
HiChIP
Hi-C
FAIRE-Seq
DNase-Seq
ChIP-Seq
CaHi-C
Bisulfite-Seq
ATAC-Seq
Aerial_rosette
Aerial_tissue
Apical_meristem
Callus
Carpel
Cotyledons
Embryo
Endosperm
Floral_bud
Flower
Flower_bud
Guard_cells
Hypocotyl
Immature_inflorescence
Inflorescence
Leaf
Meristem
N
Ovule
Plantlets
Pollen
Root
Root_hair
Root_non-hair
Root_tip
Rosette
Rosette_leaf
Seed
Seed_coat
Seedling
Shoot
Silique
Sperm_nuclei
Stem
Undifferentiated_cells
Whole_plant
Whole_rosette

Supplemental Figure 1. Distribution of data collection.

(A) Distribution of data collection by experiment type and submission date.

(B) Distribution of data collection by tissue source and mutant type.

(C) Distribution of data collection by tissue source and experiment type.

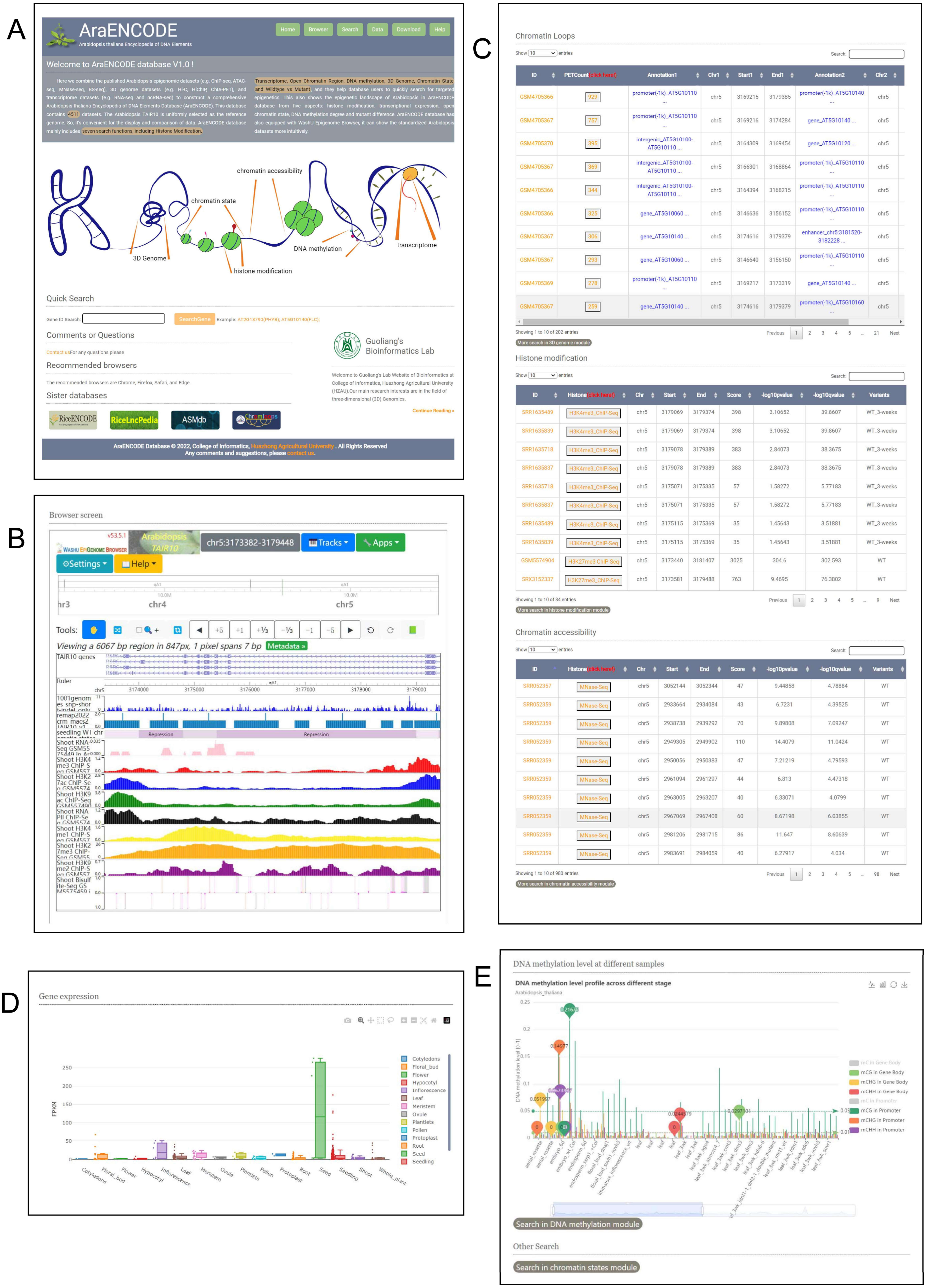
A
AraENCODE
Arabidopsis thaliana Encyclopedia of DNA Elements
Home
Browser
Search
Data
Download
Help
Welcome to AraENCODE database V1.0 !
chromatin accessibility
chromatin state
3D Genome
DNA methylation
transcriptome
histone modification
Quick Search
Comments or Questions
Recommended browsers
Sister databases
Guoliang's Bioinformatics Lab
AraENCODE Database © 2022, College of Informatics, Huazhong Agricultural University . All Rights Reserved
B
Browser screen
chr5:3173382-3179448
Tracks
Apps
Settings
Help
Tools:
Viewing a 6067 bp region in 847px, 1 pixel spans 7 bp
Metadata »
TAIR10 genes
Ruler
Repression
C
Chromatin Loops
Showing 1 to 10 of 202 entries
More search in 3D genome module
Histone modification
Showing 1 to 10 of 84 entries
More search in histone modification module
Chromatin accessibility
Showing 1 to 10 of 980 entries
More search in chromatin accessibility module
D
Gene expression
FPKM
Cotyledons
Floral_bud
Flower
Hypocotyl
Inflorescence
Leaf
Meristem
Ovule
Plantlets
Pollen
Protoplast
Root
Seed
Seedling
Shoot
Whole_plant
E
DNA methylation level at different samples
DNA methylation level profile across different stage
Arabidopsis_thaliana
DNA methylation level [0-1]
mC in Gene Body
mCG in Gene Body
mCHG in Gene Body
mCHH in Gene Body
mC in Promoter
mCG in Promoter
mCHG in Promoter
mCHH in Promoter
Search in DNA methylation module
Other Search
Search in chromatin states module

Supplemental Figure 2. An example for gene AT5G10140 (also known as FLC or RSB6) in the "Quick Search" module.

(A) AraENCODE homepage.

(B) A WashU browser window in the FLC gene region that includes a few tracks: the 7 types of histone modifications and DNA methylation as well as chromatin states and SNP information.

(C) Specific protein-mediated chromatin loops information, histone modification information, and chromatin accessibility information involved in the FLC gene.

(D) Cross-tissue differential gene expression of FLC.

(E) Differential methylation levels across samples of FLC.

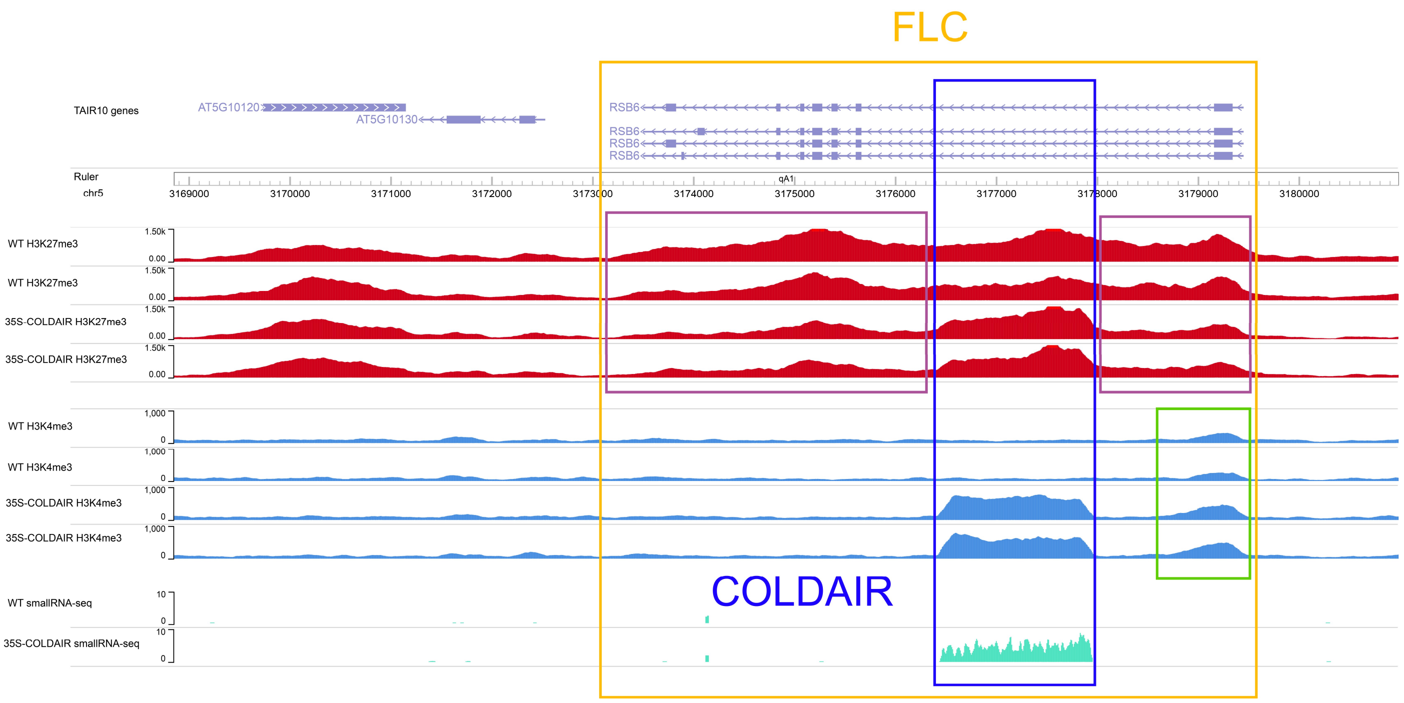


Supplemental Figure 3. The abundance levels of H3K27me3 and H3K4me3 around gene AT5G10140 (also known as FLC or RSB6) (chr5:3168852-3180985) and the expression level of long noncoding RNA COLDAIR with smallRNA-seq in AraENCODE.

The orange box is the FLC gene region. The purple boxes are genomic regions where H3K27me3 levels are significantly lower in 35S-COLDAIR mutants than in the wild type. The green box is a genomic region where H3K4me3 levels are significantly higher in the 35S-COLDAIR mutant than in the wild type.

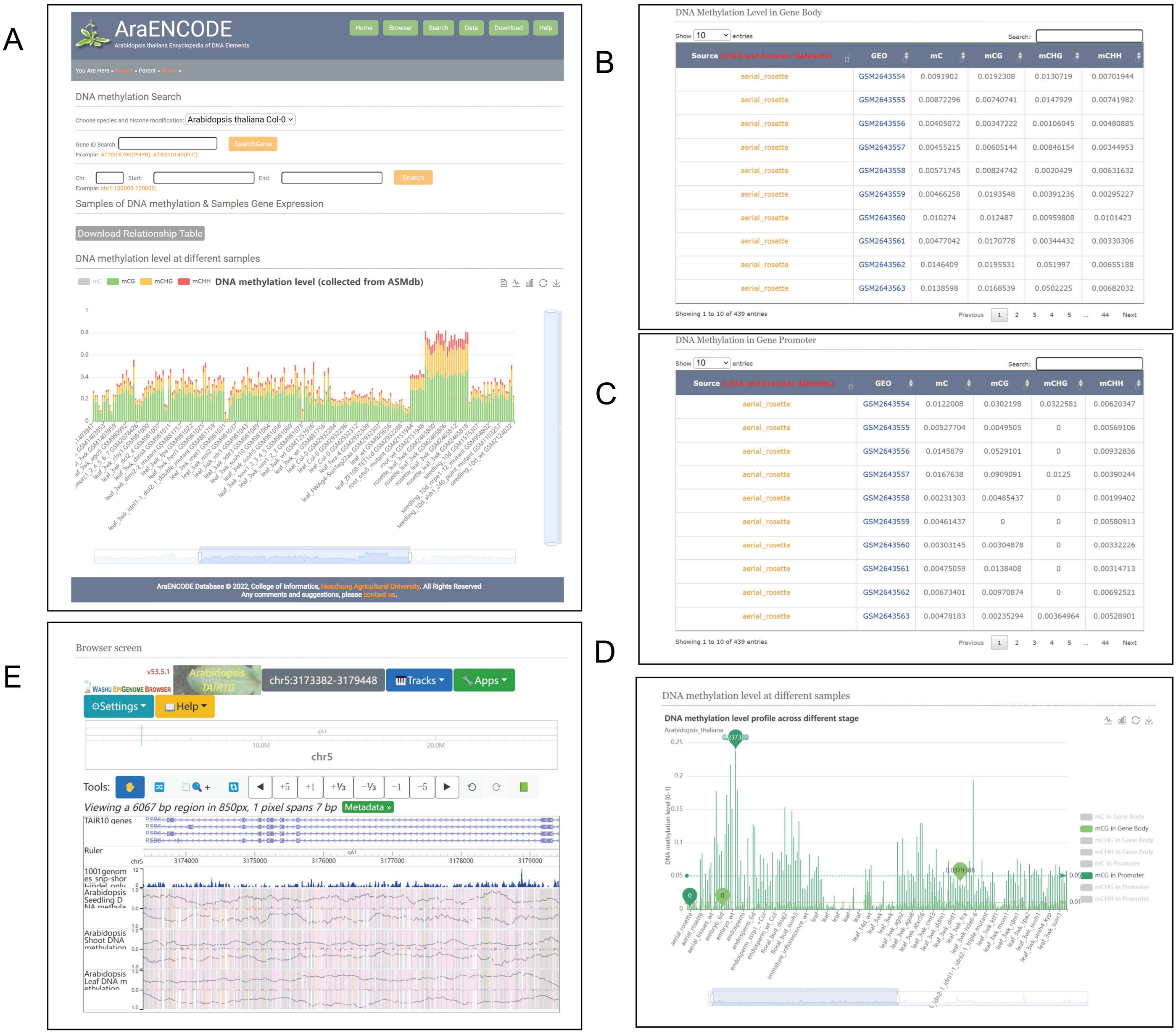


DNA Methylation Level in Gene Body

| Source (click and browser datasets) | GEO | mC | mCG | mCHG | mCHH |
|---|---|---|---|---|---|
| aerial_rosette | GSM2643554 | 0.0091902 | 0.0192308 | 0.0130719 | 0.00701944 |
| aerial_rosette | GSM2643555 | 0.00872296 | 0.00740741 | 0.0147929 | 0.00741982 |
| aerial_rosette | GSM2643556 | 0.00405072 | 0.00347222 | 0.00106045 | 0.00480885 |
| aerial_rosette | GSM2643557 | 0.00455215 | 0.00605144 | 0.00846154 | 0.00344953 |
| aerial_rosette | GSM2643558 | 0.00571745 | 0.00824742 | 0.0020429 | 0.00631632 |
| aerial_rosette | GSM2643559 | 0.00466258 | 0.0193548 | 0.00391236 | 0.00295227 |
| aerial_rosette | GSM2643560 | 0.010274 | 0.012487 | 0.00959808 | 0.0101423 |
| aerial_rosette | GSM2643561 | 0.00477042 | 0.0170778 | 0.00344432 | 0.00330306 |
| aerial_rosette | GSM2643562 | 0.0146409 | 0.0195531 | 0.051997 | 0.00655188 |
| aerial_rosette | GSM2643563 | 0.0138598 | 0.0168539 | 0.0502225 | 0.00682032 |

Showing 1 to 10 of 439 entries

DNA Methylation in Gene Promoter

| Source (click and browser datasets) | GEO | mC | mCG | mCHG | mCHH |
|---|---|---|---|---|---|
| aerial_rosette | GSM2643554 | 0.0122008 | 0.0302198 | 0.0322581 | 0.00620347 |
| aerial_rosette | GSM2643555 | 0.00527704 | 0.0049505 | 0 | 0.00569106 |
| aerial_rosette | GSM2643556 | 0.0145879 | 0.0529101 | 0 | 0.00932836 |
| aerial_rosette | GSM2643557 | 0.0167638 | 0.0909091 | 0.0125 | 0.00390244 |
| aerial_rosette | GSM2643558 | 0.00231303 | 0.00485437 | 0 | 0.00199402 |
| aerial_rosette | GSM2643559 | 0.00461437 | 0 | 0 | 0.00580913 |
| aerial_rosette | GSM2643560 | 0.00303145 | 0.00304878 | 0 | 0.00332226 |
| aerial_rosette | GSM2643561 | 0.00475059 | 0.0138408 | 0 | 0.00314713 |
| aerial_rosette | GSM2643562 | 0.00673401 | 0.00970874 | 0 | 0.00692521 |
| aerial_rosette | GSM2643563 | 0.00478183 | 0.00235294 | 0.00364964 | 0.00528901 |

Showing 1 to 10 of 439 entries

Supplemental Figure 4. An example for AT5G10140 (also known as FLC or RSB6) in the “DNA Methylation” module.
(A) AraENCODE “DNA Methylation” page.
(B) DNA methylation level in the gene body region of FLC.
(C) DNA methylation level in the gene promote region r of FLC.
(D) Differential methylation levels across samples of FLC.
(E) Browse DNA methylation landscape at single-base resolution in different samples.

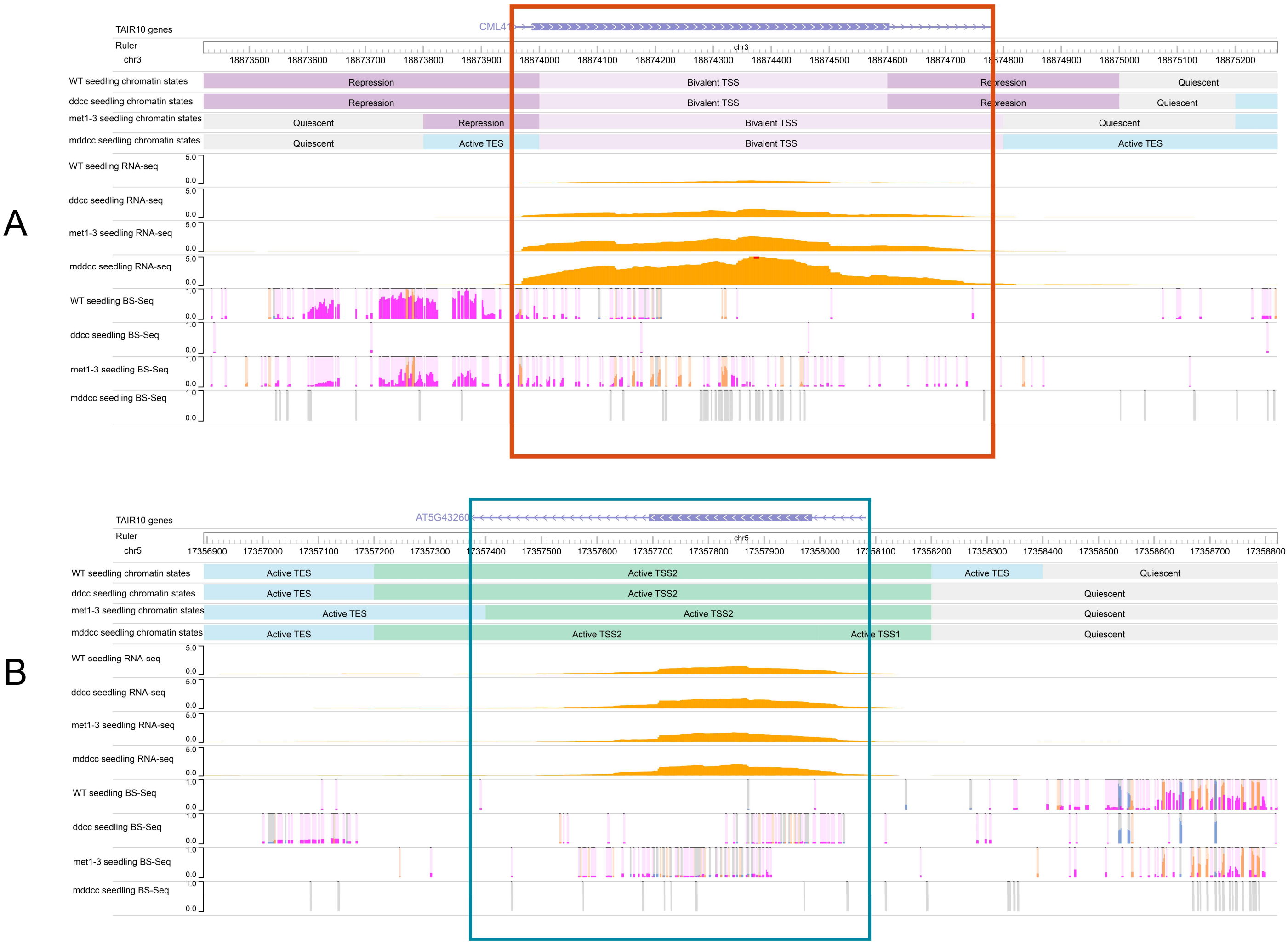


Supplemental Figure 5. The expression levels and DNA methylation levels of AT3G50770 (CML4) and AT5G43260 in wild type (Col-0) and three mutant types (non-CG methylation-free mutant met1, CG methylation-free mutant ddcc, and DNA methylation-free mutant mddcc) in AraENCODE.

(A) The expression of AT3G50770 (chr3:18873422-18875273) is up-regulated in the methylation-free mutants, especially in the mddcc mutant, whose DNA methylation in all contexts is eliminated.

(B) The expression of AT5G43260 (chr5:17356895-17358822) does not show significant differences between wild type and mutants.

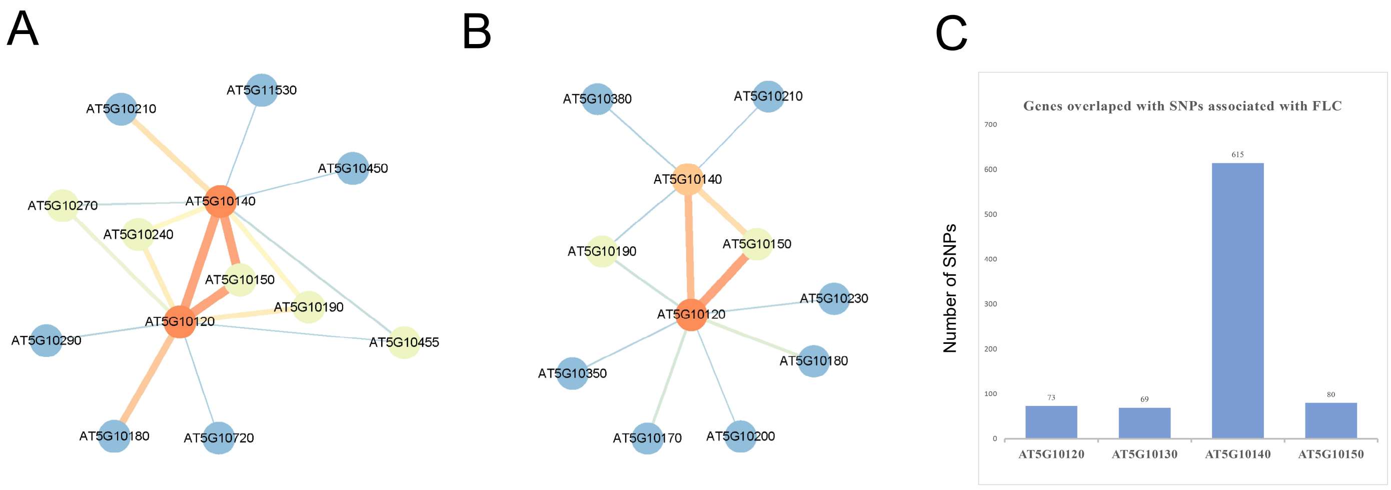


Supplemental Figure 6. The gene-gene interaction network of AT5G10140 (also known as FLC or RSB6) in AraENCODE.

(A) The gene-gene interaction network of FLC (characterized by H3K27me3 HiChIP).

(B) The gene-gene interaction network of FLC (characterized by H3K9ac HiChIP).

(C) Number of SNPs associated with FLC expression in AraGWAS, which are located in 4 different gene bodies.

### Case Study

As a well-established paradigm of epigenetic study in model plant Arabidopsis thaliana, the methyltransferase CLF (CURLY LEAF, coded by AT2G23380) and its catalyzed H3K27me3 were identified to perform biological functions, such as repressing homeotic genes and the establishment of cell identity (Goodrich et al., 1997). Characteristic phenotypes associated with mutation of CLF are upward-curled leaves with partial floral identity as well as precocious flowering (Goodrich *et al.*, 1997; Xiao et al., 2017). Reported by some studies (Huang et al., 2020; Huang et al., 2021; Rodriguez-Granados et al., 2016), histone modifications, such as H3K27me3, is linked tightly with genome structure in Arabidopsis.

To verify whether changes in histone modifications lead to alterations in chromatin structure and affect transcriptome reprogramming, ultimately changing phenotypes, we conducted a sample study using the four capture Hi-C datasets collected in AraENCODE (Huang *et al.*, 2021), which include two replicates of wild type and two replicates of clf29 mutant (Supplemental Table 3). According SCCs (stratum-adjusted correlation coefficient) between four samples, samples with the same genotype have higher similarity of chromatin interaction in Hi-C matrixes (Supplemental Figure 7A). The clf29 mutant has some specific loops which are not present in WT (Supplemental Figure 7B). These phenomena in genome structure are consistent with previous study (Huang *et al.*, 2021), showing the linkage of reversible histone modifications and 3D genome organization. Furthermore, we want to seek the particular loops associated with different genotypes, and identify whether these loops are engaged collectively in certain biological functions. These particular loops were found by hiccupsdiff, which works by checking the loops called in each hic file on the other and checks them by again calling HiCCUPS (Durand et al., 2016). We get the loops set 1 and set 2, which respectively include 187 differential loops specific to WT and 263 differential loops specific to clf29. Gene sets involved in differential loops were obtained by annotating genes to anchors based on genomic location. Gene set 1 and 2 were used to make Gene Ontology (GO) enrichment analysis respectively. Important biological process were determined, such as flowering development, which could lead to early flowering phenotype in clf29 (Supplemental Figure 7C,D). Furthermore, these 39 genes involved in flowering development are distributed throughout the whole genome in Arabidopsis (Supplemental Figure 7E). Hence, there is little likelihood that these genes are captured together due to their proximity on the genome. Among these genes, AT2G06050 (ATOPR3) and AT5G65080 (MAF5) show significant upregulation in expression levels (p-value<0.05 and fold change) (Supplemental Figure 7F). MAF5, like other MAF genes (e.g. FLC), is a repressor of flowering and is upregulated during vernalization (Ratcliffe et al., 2003; Sheldon et al., 2009). In clf29 mutant, MAF5 showed stronger interaction with some distant enhancers than WT, which may be involved in the upregulation of the MAF5 (Supplemental Figure 7G).

In this case study, we integrated data to check the differences of chromatin conformation between clf29 and WT, and found that some genes which are located on these regions with differential structure are involved in the same biological process and act on specific phenotypes. We found that a flowering-related gene, MAF5, has upregulated expression level and strong interaction strength with surrounding the enhancers in clf29. We believe that such information can provide new insights for future research and our database can facilitate researchers to use the existing data.

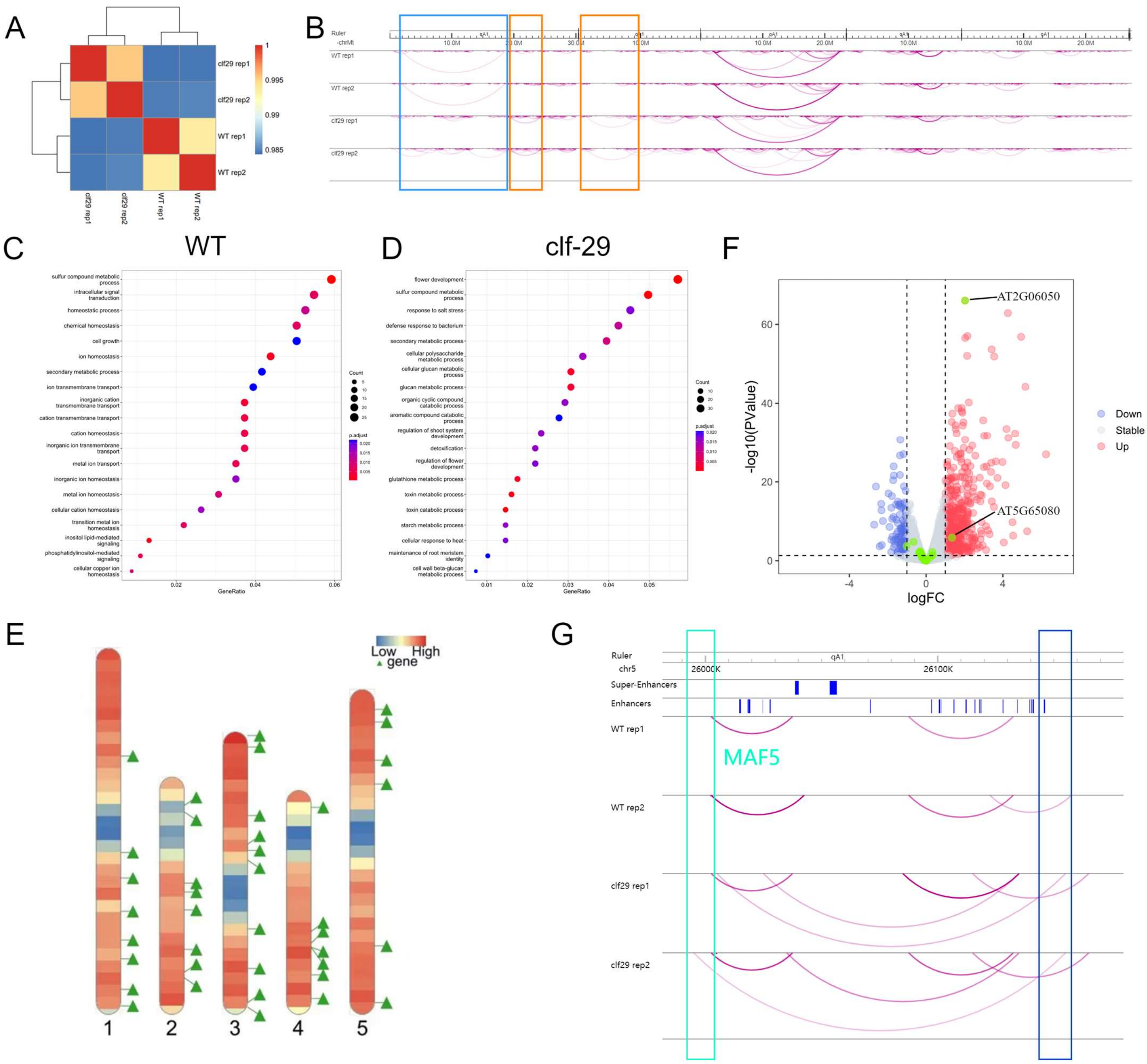


Supplemental Figure 7. A case study for AraENCODE.

(A) SCCs (stratum-adjusted correlation coefficient) between four samples. samples with same genotype have higher similarity of chromatin interaction in CaHi-C matrixes

(B) Loops detected by HiCCUPS on interaction matrixes. The orange boxes are clf29-specific loops. The blue box is a WT-specific loop.

(C) Gene Ontology (GO) enrichment analysis using genes overlapped with differential loops' anchors of WT.

(D) Gene Ontology (GO) enrichment analysis using genes overlapped with differential loops' anchors of clf29.

(E) Distribution of 39 genes, which are involved in flowering development and are overlapped with clf29 specific loops.

(F) Differential gene expression of clf29 and WT. 39 Genes related to flowering development are marked as green. AT2G06050 (ATOPR3) and AT5G65080 (MAF5) show significant upregulation in expression levels (p-value<0.05 and fold change).

(G) Different interaction strength of MAF5 with surrounding enhancers in clf29 and WT.

**Supplemental Table 3. Sample information.**

| GEO | Run ID | Ecotype | Library | Genotype | Tissue |
|---|---|---|---|---|---|
| GSM4705372 | SRR12361554 | Col-0 | CaHi-C | WT | Shoot |
| GSM4705374 | SRR12361556 | Col-0 | CaHi-C | WT | Shoot |
| GSM4705375 | SRR12361557 | Col-0 | CaHi-C | clf29 | Shoot |
| GSM4705376 | SRR12361558 | Col-0 | CaHi-C | clf29 | Shoot |
